# Multimodule imaging of the hierarchical equine hoof wall porosity and structure

**DOI:** 10.1101/2023.06.21.545722

**Authors:** Mahmoud A. Mahrous, Charul Chadha, Pei L. Robins, Christian Bonney, Kingsley A. Boateng, Marc Meyers, Iwona Jasiuk

## Abstract

The equine hoof wall has a complex, hierarchical structure that can inspire designs of impact-resistant materials. In this study, we utilized micro-computed tomography (μ-CT) and serial block-face scanning electron microscopy (SBF-SEM) to image the microstructure and nanostructure of the hoof wall. We quantified the morphology of tubular medullary cavities by measuring equivalent diameter, surface area, volume, and sphericity. High-resolution μ-CT revealed that tubules are partially or fully filled with tissue near the exterior surface and become progressively empty towards the inner part of the hoof wall. Thin bridges were detected within the medullary cavity, starting in the middle section of the hoof wall and increasing in density and thickness towards the inner part. Porosity was measured using three-dimensional (3D) μ-CT, two-dimensional (2D) μ-CT, and a helium pycnometer, with the highest porosity obtained using the helium pycnometer (8.07%), followed by 3D (3.47%) and 2D (2.98%) μ-CT. SBF-SEM captured the 3D structure of the hoof wall at the nanoscale, showing that the tubule wall is not solid, but has nano-sized pores, which explains the higher porosity obtained using the helium pycnometer. The results of this investigation provide morphological information on the hoof wall for the future development of hoof-inspired materials and offer a novel perspective on how various measurement methods can influence the quantification of porosity.

## 1. Introduction

The remarkable properties of biological materials, including self-healing, adhesion, and impact resistance, offer inspiration for developing advanced engineering materials [1]. Impact resistance refers to a material’s ability to withstand intense forces or shocks. Porosity is a common feature of both naturally occurring impact-resistant materials and those used for engineering applications in biomedical, aerospace, automotive, and packaging industries [2-5]. The hoof wall is a porous material responsible for protecting the internal structure of the hoof from impact forces generated during contact with the ground at high speed.

The hoof wall has a complex, hierarchical structure, shown in Fig. 1, providing high impact resistance and fracture toughness [6, 7]. At the nano-scale, intermediate filaments (IFs) (∼7-10nm) act as fibers embedded in an amorphous protein matrix [8]. Aligned IFs form macrofibrils, roughly 700 nm in diameter that are dispersed inside disk-shaped cells around 10-40 μm across and 5 μm thick [9]. Concentric lamellae, each made from a single layer of cells, create cylindrical structures called tubules that run from the top to the bottom of the hoof wall [10, 11]. The tubules have a 200-300 μm diameter, with a central medulla, or tubule medullary cavity (TMC), of about 50 μm [6, 12]. Intertubular regions consist of lamellae at an oblique angle with the long axis of the tubules [6, 10, 11, 13].

**Fig. 1.**
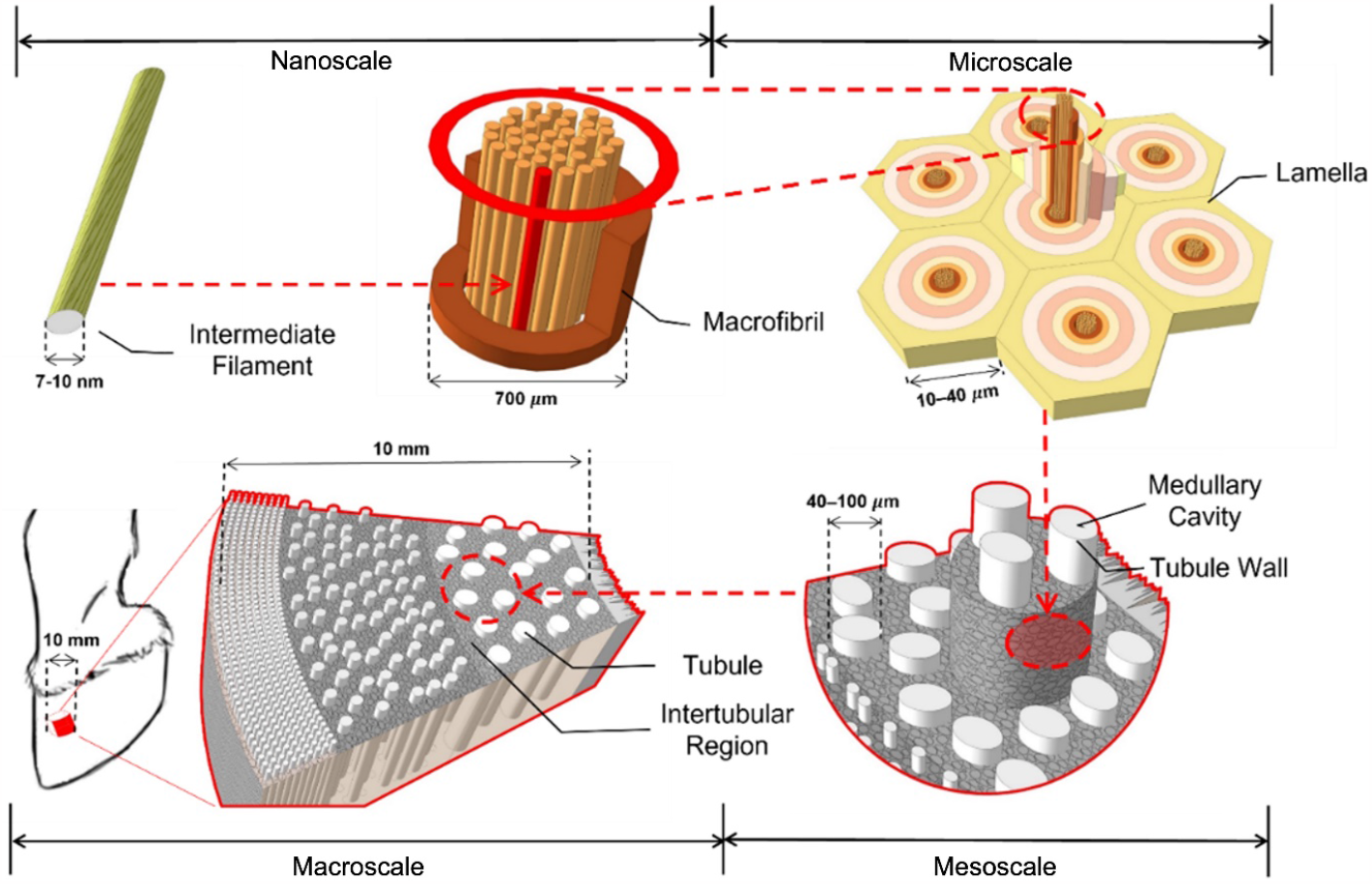
The hierarchical structure of an equine hoof wall

Figure 2 depicts the coordinate axes typically used to describe locations in the hoof wall. The tubules’ axes are parallel to the outer surface, which defines the longitudinal direction. Orthogonal to the longitudinal axis is the radial direction that describes locations along the thickness [6, 12]. Most of the hoof wall is the stratum medium except for thin layers on the outer and inner surfaces referred to as the stratum externum and stratum internum, respectively [14]. The hoof wall is further divided along its circumferential or transverse direction into two side regions (medial and lateral) and a front-facing (toe) region. Studies have shown that the tubule density and shape are not constant throughout the hoof wall [6, 15]. Lancaster *et al*. [16] counted the tubules and calculated their density, finding a change along the radial direction and a significant difference between the lateral, medial, and toe regions. Most research has focused on the stratum medium of the toe for consistency due to heterogeneity throughout the macroscale structure.

**Fig. 2.**
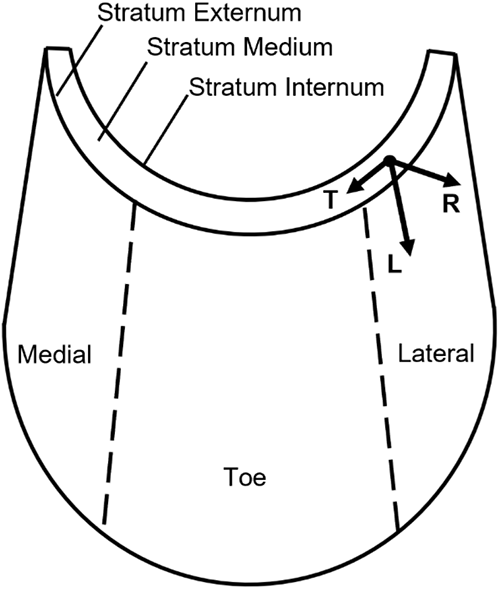
Coordinate axes and locations of the equine hoof wall

Previous studies have used the histological examination technique to study the hoof wall structure, which involved sectioning a small strip for polarized or optical microscopy [6, 17]. This process is both challenging and time-consuming and can damage the tubules’ microstructure during sample preparation. Also, histological examination is limited to a small number of tubules. Micro-computed tomography (μ-CT) allows for the scanning of larger areas at micro-level resolution and has proven to be a valuable technique for visualizing the intricate inner structures of biological materials [18-23]. Huang *et al*. [9] analyzed the structure of the hoof wall with μ-CT to find the average area fraction (∼ 3%,) and diameter (41± 9 μm) of the TMC. Then, Lazarus *et al*. [24] showed, using μ-CT, that the tubules are not continuous hollow structures but are segmented by bridges. They reported the average bridge width (10.3 ± 2.4 μm), bridge density (0.009 ± 0.002 bridges/μm), tubule density (15.91 ± 0.4 tubules/mm^2^), and porosity (0.77 ± 0.3%). However, no work was done to visualize and quantify nanoscale porosity using a 3D method. In this study, we quantified the hoof wall structure and porosity by utilizing high-resolution characterization techniques, including μ-CT, scanning electron microscopy (SEM), and serial block face scanning electron microscopy (SBF-SEM), coupled with a helium pycnometer, to measure open porosity.

SBF-SEM is a volume scanning electron microscopy that produces 3D images at nanoscale resolution (∼ 5 × 5 × 5 nm^3^) [25]. This technology is like an automated serial transmission electron microscope and requires specialized sample preparation methods, including staining the sample with various solutions, to improve image resolution. SBF-SEM is widely used to study plant and cell tissue nanostructures [26-30] but it has not been utilized for hoof wall characterization. Previously, high-resolution transmission microscopy (TEM) was used to visualize cell boundaries in the hoof wall [9]. TEM can image serial sections of a millimeter-sized specimen with a Z-resolution between 1 and 50 nm [31]. Still, there is a risk of losing sections during sample preparation, which has not been reported with SBF-SEM. Also, the more complicated sample preparation for TEM can damage the microstructure of the hoof wall tissue, while SBF-SEM prevents such damage. On the other hand, SBF-SEM produces large datasets that make reconstructing a 3D volume computationally expensive and time-consuming. Furthermore, the limited sample size of a few micrometers in the X, Y, and Z directions adds another constraint to using SBF-SEM as a characterization technique [32, 33]. In this paper, we utilized SBF-SEM to capture features of the hoof wall nanostructure, including the cell structure and the intertubular region, which would be difficult to obtain with other 2D techniques.

Hoof wall samples were first scanned using low-resolution μ-CT (13.35 μm) to visualize and quantify the tubules. Smaller samples were then extracted to study the inner section of the tubules (TMC) using higher resolution μ-CT (0.53 μm), followed by SBF-SEM to obtain a 3D block image for inner and outer hoof wall tubules as well as the intertubular region at the nanoscale. Open porosity was determined using a helium pycnometer. It will be shown that integrating these advanced techniques enhances our understanding of keratin-based materials and clarifies the effect of measurement techniques on quantifying porosity.

The paper consists of four sections and an Appendix. Section 2 describes methods, Section 3 presents results and their discussion, while Section 4 gives conclusions and future directions. The Appendix shows extra details on μ-CT and SBF-SEM imaging.

## 2. Materials and Methods

The Veterinary Department of the University of California at Davis supplied horse hooves from a mature racehorse of unknown age and sex. The hooves were removed, refrigerated within 48 hours, and then frozen at -20^°^. After thawing, hooves were cut down to isolate the hoof wall, and samples were extracted from the toe region with a bandsaw.

### 2.1 X-ray microcomputed tomography (μ-CT)

Four 3×1×1 cm^3^ samples were collected from various locations in the toe region for a comprehensive statistical analysis of the TMCs. Scans were performed using a North Star Imaging X5000 instrument (North Star Imaging, Rogers, MN, USA). Samples were scanned using a voltage of 90kV and a current of 150 μA by an X-ray source (XRayWorX [P19-701]), which has a 13.5 μm focal spot size. Multiple resolutions were evaluated to ascertain the optimal resolution for capturing tubular features. Ultimately, a maximum voxel size of 13.35 μm was determined to be the most suitable. To achieve the desired resolution and sharpness, the distances of the detector and object from the source were 975.478 and 102.545 mm.

For higher resolution imaging with a pixel size of 0.53 μm, cubic samples with a size of 2×2×2 mm^3^ were extracted from three different locations throughout the thickness of the hoof wall, as depicted by the red cubes in Fig. 6a. The scans were performed using the high-resolution Rigaku Nano 3Dx computed tomography instrument (Rigaku, Tokyo, Japan). The system is equipped with three different lenses that can be adjusted to achieve varying image resolutions, an X-ray detector, and a Peltier-cooled CCD camera capable of producing images up to 3200×2400 pixels. In this case, a resolution of 0.53 μm was achieved by using a lens with a field of view of 3.5× 2.6 mm. The samples were scanned using a quasi-monochromatic X-ray source, with a voltage of 130 kV and a current of 61 μA produced by a Copper target.

### 2.2 Helium pycnometer

The helium pycnometer was used to determine the open porosity of the hoof wall. Six specimens, each with a dimension of 1×1×0.9 cm^3^, were used to measure the open porosity through the weighing method. The samples were immersed in water for 24 hours to ensure that all the pores were fully saturated. The samples were then removed from the water, and the outer surface was meticulously cleaned to eliminate any residual water and ensure that only the inner pores of the sample were filled. Samples were weighed precisely using an analytical balance with an accuracy of ±0.0001g to obtain the bulk mass. The volume was calculated using the pycnometer instead of a standard caliber, as the samples have some irregularity on the outer surface. The samples were individually placed and measured in the AccuPyc 1330 instrument (Micromeritics Instrument Corp., Norcross, GA, USA). This instrument uses pressurized helium gas, as it is a monomolecular gas with a diameter of around 0.22 nm, smaller than the diameter of water vapor (0.28 nm), allowing for the detection of nanoscale pores. The instrument measures the volume of the sample, and using the known weight (measured previously), the bulk density of each sample was calculated. The samples were then removed from the pycnometer and placed in an oven at 80°C for 24 hours to remove the water from the pores of the hoof wall. This temperature was carefully selected after several trials to prevent the over-drying of the sample. Over-drying could lead to the formation of cracks in the sample structure, resulting in erroneous porosity results. The dried samples were weighed to obtain the mass and then placed back into the helium pycnometer to measure the sample volume and true density in a dry state. This approach enabled accurate measurement of the amount of helium gas that replaced the air in the inner pores of the sample. By knowing the bulk density of the sample and its true density, the sample porosity was calculated using the following equation [34]:

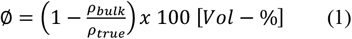

where ∅ is the percent porosity, *ρ*_*bulk*_ is the density of the wet sample, and *ρ*_*true*_ is the density of the dried sample.

### 2.3 Scanning electron microscopy (SEM)

For SEM scanning, a 2 × 2 × 2 mm^3^ sample was removed from location C in Fig. 6a. The sample was mounted on a metal holder and coated with a thin layer of 5 nm gold-palladium in a low-vacuum Emscope SC 500 Sputter Coater. The coating helps to reduce thermal damage to the hoof wall tissue and minimize the charging effect on the sample surface during the scan. The sample was analyzed using high vacuum mode on an Axia ChemiSEM instrument (Thermo Fisher Scientific, Cleveland, OH, USA) with an acceleration voltage of 30 kV.

### 2.4 Serial Block-Face Scanning Electron Microscopy (SBF-SEM)

A sample with dimensions of 2×2×3 mm^3^ was embedded in epoxy using the standard serial block-face SEM protocol [33,34] with modifications to extend the time. The sample was then treated with three solutions for different durations: 2% osmium in 1.5% potassium ferrocyanide for three days, 0.01% thiocarbohydrazide (TCH) solution for one day, and 2% osmium for one day. The sample was then dehydrated and embedded in Durcupan. The epoxy-embedded samples were mounted on aluminum pins (Gatan) using silver epoxy (Ted Pella, Redding CA) and sputter-coated with a thin layer of Au/Pd before being subjected to block-face imaging. Serial block-face imaging was performed using a Sigma VP (Zeiss, Oberkochen, Germany) equipped with a Gatan 3View system (model: 3View2XP) and a nitrogen gas injection manifold (Zeiss model 346061-9002-200). For this work, the samples were typically imaged at 2.0 keV, with 50 nm cutting intervals, a 1.0 nm pixel size (12k × 12k pixels), a beam dwell time of 1.0 μsec, and a high vacuum chamber pressure of approximately 5×10^−3^ mbar.

## 3. Results and discussion

### 3.1 Statistical analysis

A total of 3905 tubules (tubules referred to here as the TMCs region, not including the tubule wall) were segmented and isolated from the hoof wall matrix using the segmentation and labeling process described in the Appendix. They were analyzed to obtain the area, volume, sphericity, and equivalent diameter (Eqdiameter) of the TMCs. Eqdiameter is calculated using the following equation [38]:

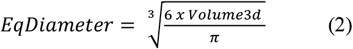

The results of the statistical analysis obtained from the segmented 3D μ-CT scans are summarized in Table 1. The analysis includes the mean, median, and standard deviation (Std. Dev). Figure 3a exhibits the average frequency distribution of TMC Eqdiameter. The results indicate that the Eqdiameter of the TMCs ranges from 55 μm to 153.16 μm, with an average of 69.43 μm. Most TMCs have an Eqdiameter ranging from 55.91 μm to 75.91 μm, accounting for 48.8% of the tubules analyzed in this study. Only 2.8% of the TMCs have an Eqdiameter greater than 110 μm.

**Table 1.** Summary of statistical results for different geometric descriptors of TMCs in the hoof wall

| Geometric Descriptor | Mean | Median | Std. Dev. |
| --- | --- | --- | --- |
| Eqdiameter ($\mu\text{m}$) | 69.43 | 66.66 | 16.36 |
| Surface area ($\mu\text{m}^2$) | $3.55 \times 10^4$ | $2.44 \times 10^4$ | $3.45 \times 10^4$ |
| Volume ($\mu\text{m}^3$) | $4.53 \times 10^4$ | $2.85 \times 10^4$ | $4.93 \times 10^4$ |
| Sphericity | 0.86 | 0.86 | 0.09 |

**Fig. 3.**
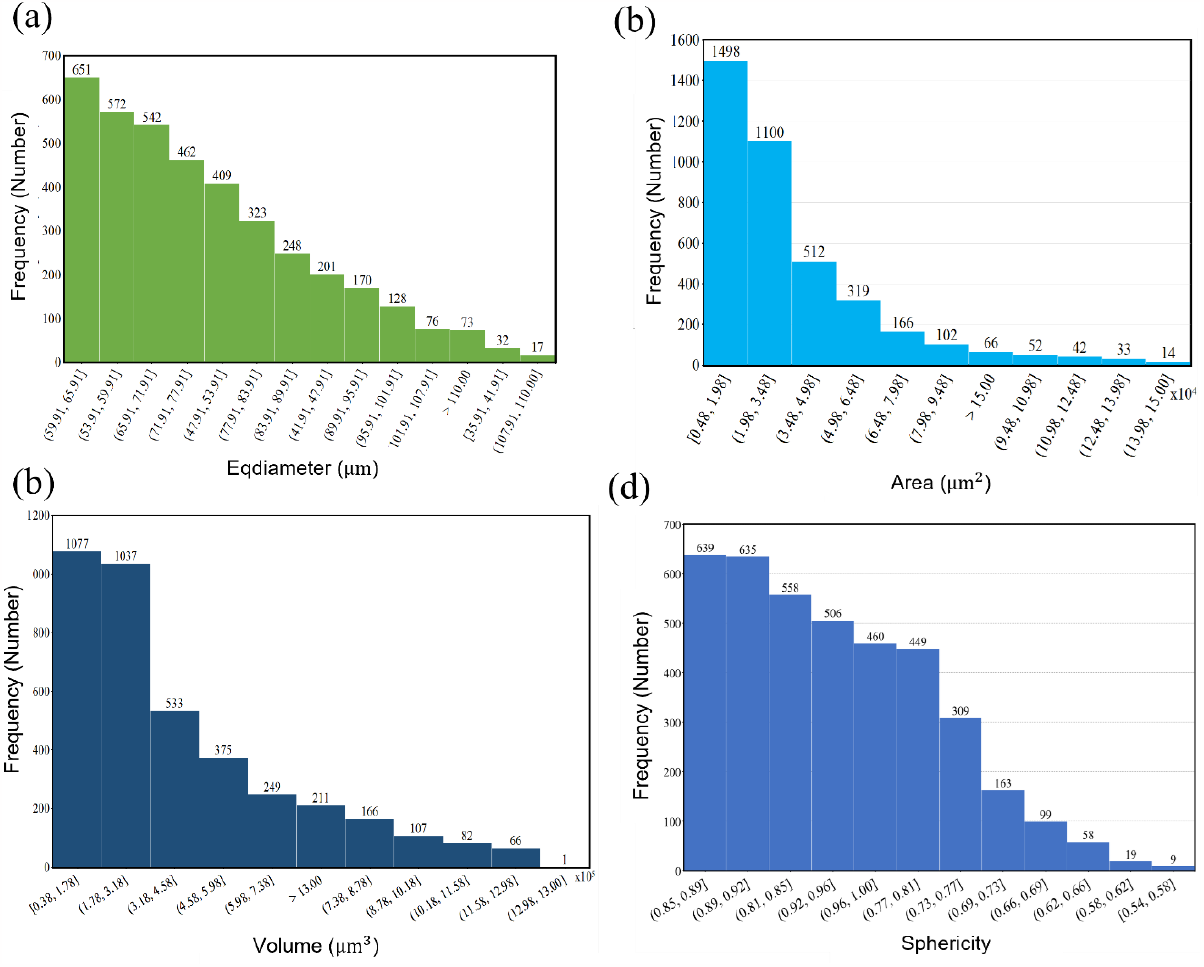
(a) Histogram of the Eqdiameter size distribution. (b) Histogram of TMC’s area size distribution. (c) Histogram of TMC’s volume size distribution. (d) Histogram of TMC’s sphericity size distribution.

Previous studies have reported different TMC Eqdiameter values, including 41 ± 9 micrometers (Huang *et al*. [9]), 200 μm (Kasapi *et al*. [35]), and 300 μm (Kasapi *et al*. [36]). The significant differences between these results and those of our study are likely due to the location of the sample within the hoof and the analysis techniques used. Our results, obtained using the 3D μ-CT technique, minimize the impact of sample location by covering multiple areas within the hoof wall. Additionally, our analysis was based on approximately 1500 images.

Figure 3b shows the distribution of TMC surface area measurements. The analysis reveals that the surface area of 2598 tubules (66.5%) ranges from 0.48 × 10^4^ μm^2^ to 3.48 × 10^4^ μm^2^, making it the dominant size. The second largest surface area, representing 25.5%, ranges from 3.48 × 10^4^ μm^2^ to 7.98 × 10^4^ μm^2^. The remaining TMC surface areas range from 7.98 × 10^4^ μm^2^ to 15 × 10^4^ μm^2^, with 66 TMCs measuring more than 15 × 10^4^ μm^2^, having a maximum surface area of 56.48 × 10^4^ μm^2^. Additionally, a calculation was performed to determine the sample’s porosity based on the total surface area of the TMCs relative to the total surface area of the sample using the following formula.

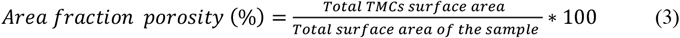

Using Equation 3, we obtained a porosity value of 2.98%, close to the previously recorded value of 3% by Huang *et al*. [9]. Our results are based on the analysis of a large sample size of 3905 tubules, which provides enough confidence in the accuracy of the obtained porosity value. In addition, previous work, which dealt with the hoof wall porosity calculation, was done by counting the number of tubules relative to the sample area, using primarily light [15, 17] and electron microscopy techniques [37]. This method can be prone to human error and can only provide limited information about a specific location which may not accurately represent the entire hoof wall porosity.

We used an automated μ-CT method to eliminate the possibility of human error and obtain more accurate results. The only drawback of using μ-CT is identifying the best-fit threshold values that cover the entire range of densities within the sample. This identification was not difficult in our case as the hoof wall consists of two distinct regions, the TMC region (lower density) and the intertubular region (higher density), which can be easily separated using the interactive threshold modules in the Amira software. The details of this process are discussed in the μ-CT segmentation section of the Appendix.

The volume size distribution of the TMC is depicted in Fig. 3c. The volume of the TMC varies from 0.38 × 10^5^ μm^3^ to 73.63 × 10^5^ μm^3^. Most volumes are between 0.38 × 10^5^ μm^3^ and 4.58 × 10^5^ μm^3^, accounting for 67.78% of the total analyzed TMCs. Additionally, 5.4% of the TMCs (211 TMCs) have a volume greater than 13 × 10^5^ μm^3^. As shown in Fig. 3c, there is a gradual decrease in the TMC volume until it reaches the minimum volume of 13 × 10^5^ μm^3^, indicating a significant variation in the TMC volume. The following equation was used to calculate the volumetric porosity of the hoof.

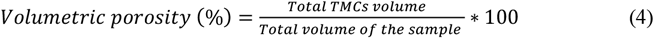

The porosity calculated using the 3D volume method is 3.47%, slightly higher than the porosity calculated using the 2D surface area method. The 2D analysis assumes that the pores are perfectly circular, neglecting the irregularity of the pore walls, which could result in inaccurate results. Chandrappa *et al*. [38] concluded that two pores with similar cross-sectional areas would have equal diameters, even with different shapes, surface areas, and volumes. Thus, the 3D pore parameters are more accurate for porosity analysis, as the analysis is based on the voxel size, not the radius of the cavity, unlike the 2D image analysis method.

Sphericity, *ψ*, is a dimensionless parameter used to assess the degree of the spherical shape of an object. It is frequently used to quantify the shape of pores in biological materials [39]. Determining the particle sphericity with high accuracy is challenging as it requires precise measurements of the particle’s surface area and volume in three dimensions [40]. The calculation of sphericity is done by dividing the surface area of an equal-volume sphere by the actual surface area of the object (as per equation 5). Sphericity values range from 0 to 1, with 1 indicating a perfect spherical shape [41]:

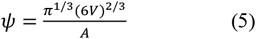

where *V* is the volume of the TMC and *A* is the TMC surface area [42].

The frequency distribution of the sphericity measurements of the TMCs is demonstrated in Fig. 3d. The results indicate that the sphericity of the pores ranges from 0.54 to 1, with most of them having a sphericity value of 0.73 to 0.92 (65.71%). Table 1 shows that the mean sphericity is 0.86, suggesting that most TMCs are not perfect spheres or entirely elliptical. Huang *et al*. and Kasapi *et al*. [6, 9] described the sphericity of the TMCs based on the lengths of the major and minor axes without giving a well-defined value for the sphericity.

Figure 4 and Table 2 present the results of the area fraction porosity obtained from low-resolution (13.35 μm) μ-CT images. The overall area fraction porosity for the entire hoof wall sample is approximately 3.88%, slightly higher than the value reported by Huang *et al*. [9]. The results indicate that the average area fraction porosity increases from Zone 1 (the inner zone) to Zone 4 (the outer zone) of the hoof wall. However, Zone 3 showed a lower area fraction porosity than Zone 2 but higher than Zone 1. The results reported by Reilly *et al*. [15] indicate that tubule density increases as the zone moves from the inner to the outer equine hoof wall. This difference between our work and Reilly *et al*. [15] arises because we partitioned the hoof wall differently. Reilly *et al*. [15] divided the stratum medium of the hoof wall into four zones at 25%, 47%, and 69% of the hoof wall depth. In our work, we included the stratum medium and part of the stratum externum, and divided the hoof wall into four equal zones at 25%, 50%, and 75% of the hoof wall depth.

**Table 2.** A summary of area fraction porosity (%) results for different zones.

| Zone | Mean | Median | Std. Dev. |
| --- | --- | --- | --- |
| 1 | 3.49 | 2.87 | 1.23 |
| 2 | 4.07 | 3.60 | 1.42 |
| 3 | 3.65 | 2.83 | 1.62 |
| 4 | 4.21 | 4.60 | 1.10 |

**Fig. 4.**
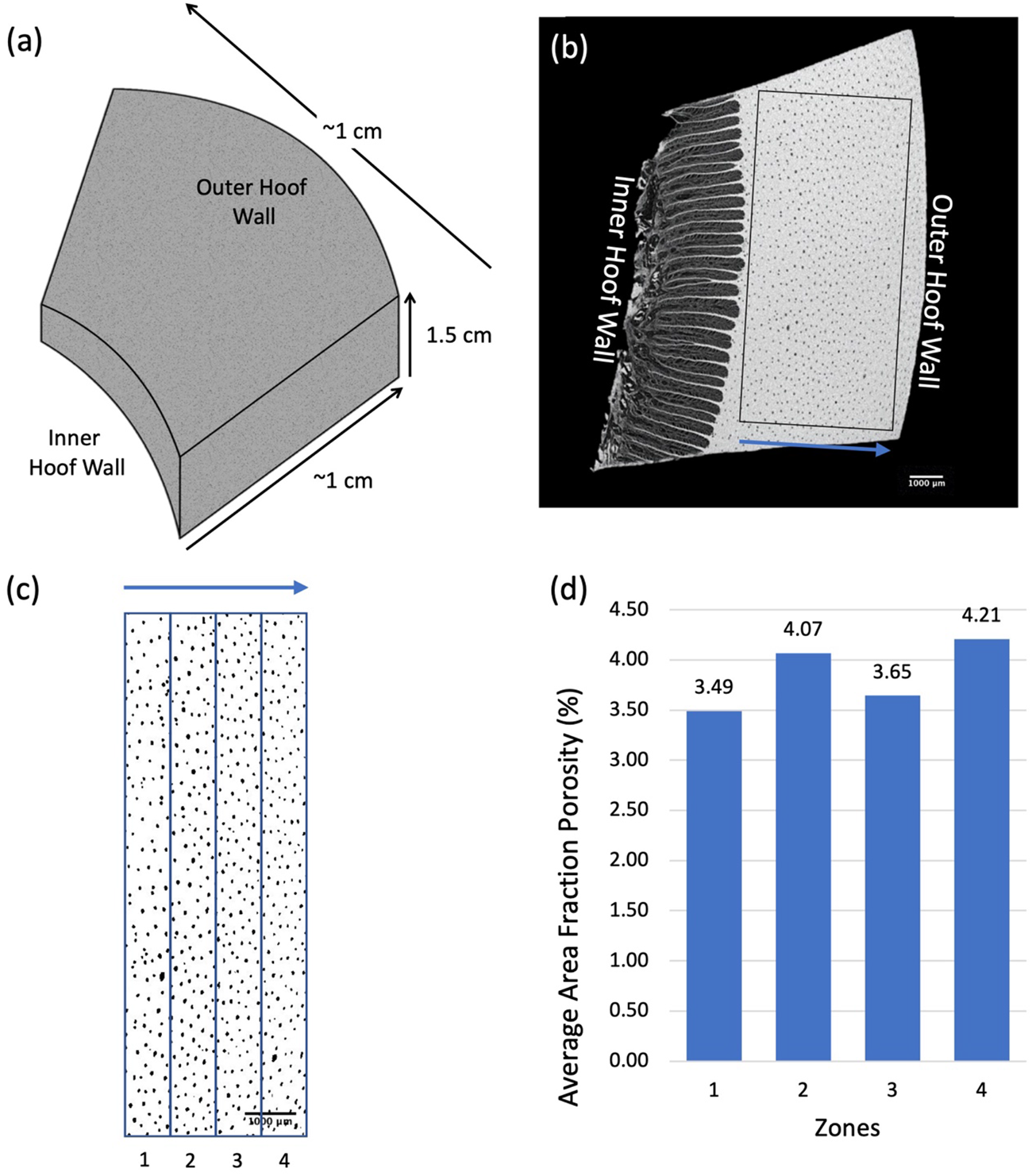
Flowchart for the image analysis process used to determine the area fraction porosity percentage. The chart includes the following steps: (a) 3D reconstruction of the μ-CT scan, (b) selection of a random image for analysis, where the black box represents the cropped area, (c) division of the cropped area into 4 zones at 25%, 50%, and 75% of the hoof wall depth, and (d) a histogram showing the results of the area fraction porosity, which is averaged for each zone.

### 3.2 Low-resolution μ-CT

Figure 5a presents a μ-CT image of a 3D volume rendering of a hoof wall sample scanned with a 13.35μm voxel size resolution. The image illustrates that the tubules are well-aligned and run parallel to each other. The top view of the 3D reconstructed image of the segmented TMC (Fig. 5b and 5c) reveals that the tubules are arranged parallel to the longitudinal direction and randomly distributed in the radial direction. Figure 5d shows a waviness in the tubular structure, which was previously reported by Lazarus *et al*. [43]. This waviness might be caused by the dryness and loss of moisture from the hoof sample during the micro-CT scanning, which can result from the heat generated from the X-ray beam. Further research could assess the impact of waviness on hoof wall’s mechanical properties and its energy absorption capacity. Figure 5e shows that TMCs are segmented into air pockets separated by thin tissue (bridges), similar to the findings of Huang *et al*. [9] and Lazarus *et al*. [24]. The air pockets do not have the same size and shape.

**Fig. 5.**
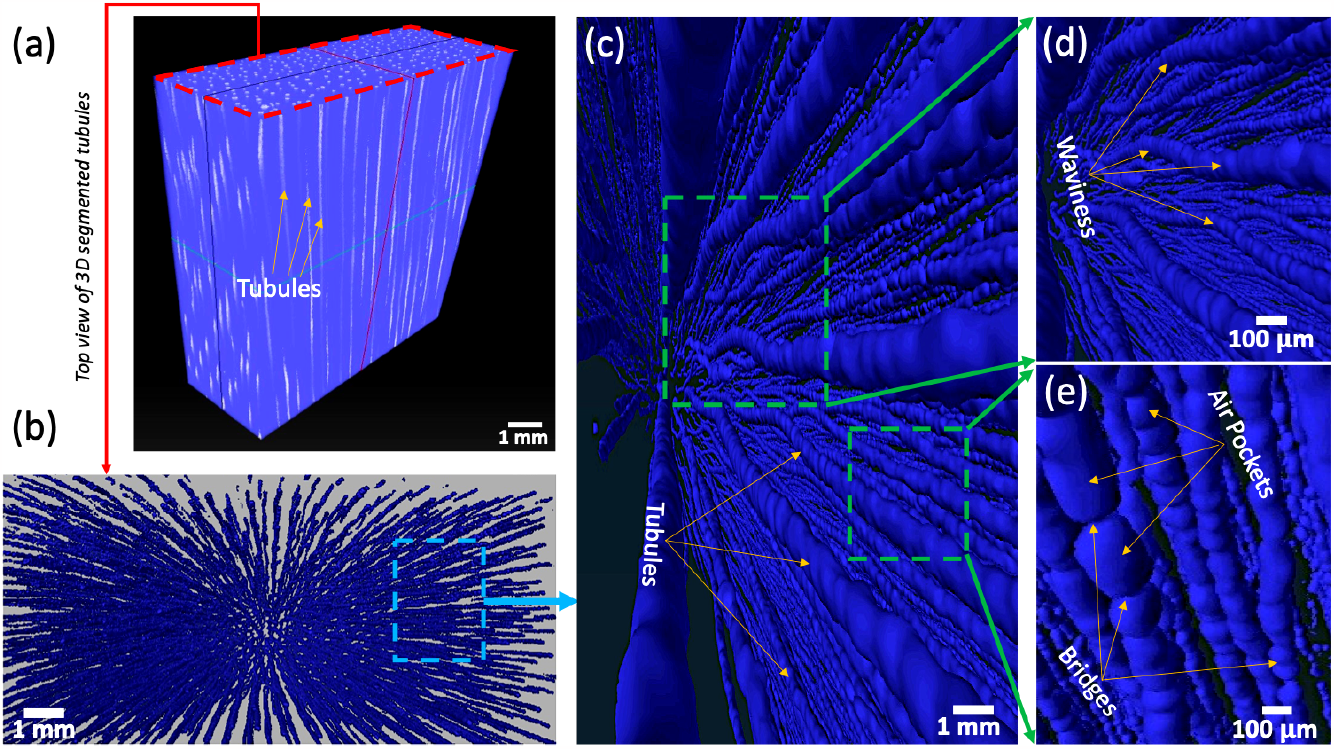
3D X-ray Micro-CT visualization of hoof wall specimen a) 3D reconstruction of hoof wall sample with a voxel size resolution of 13.35 μm, b) top view of 3D segmented (TMCs), c) longitudinal view of TMCs demonstrating their parallel alignment, d) magnified view of tubules exhibiting periodic waviness, and e) magnified view of TMCs highlighting the discontinuity of the modular cavity and the presence of bridges.

### 3.3 High resolution μ-CT

To better understand the inner structure of the TMC, high-resolution μ-CT with a resolution of 0.53μm was performed on samples extracted from three different radial locations of the hoof wall, as shown in Fig. 6a. The μ-CT images shown in Fig. 6 include axial, coronal multiplanar reconstructed, and sagittal images. In Fig. 6b, an axial view of a μ-CT image from location A (the outermost layer or dorsal region of the hoof wall) reveals that some of the TMC regions at this location are closed and fully filled with tissue. In contrast, others are partially filled with soft tissue, contrary to previous beliefs that they were empty. The coronal multiplanar reconstructed image in Fig. 6e also indicates that the TMCs are not hollow but filled with tissue in the outer section of the hoof wall.

**Fig. 6.**
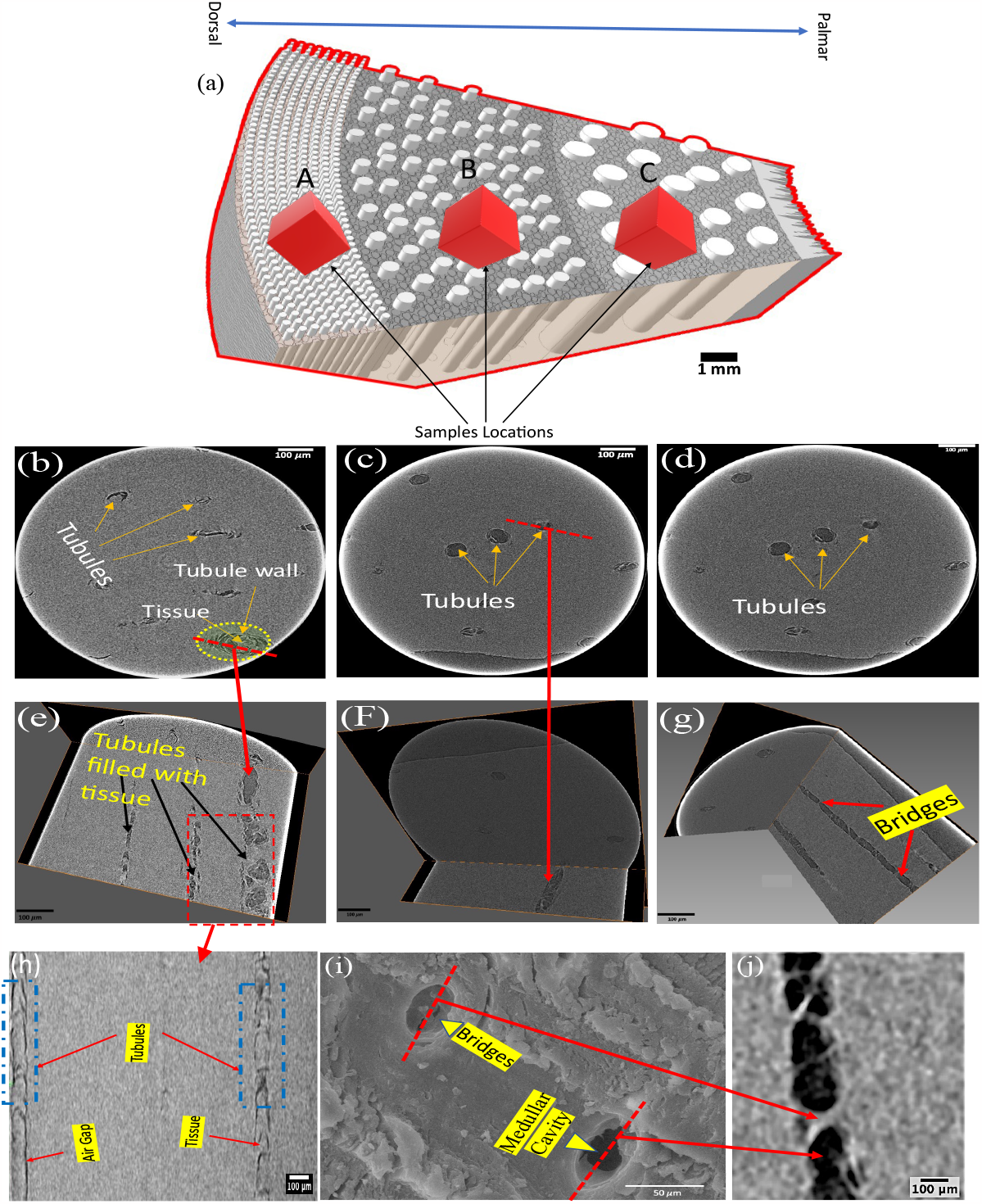
High-resolution X-ray micro-computed tomography (μ-CT) images of a hoof wall specimen. μ-CT images obtained at a resolution of 0.53 μm, with a) a schematic representation of the hoof wall structure indicating the locations of the examined samples, b) an axial view of a μ-CT image from location A (the outermost region), c) an axial view from location B, d) an axial view from location C, e) a coronal multiplanar reconstructed image of the tubules at location A, f) a coronal multiplanar reconstructed image of the tubules at location B, g) a coronal multiplanar reconstructed image of the tubules at location C, h) a coronal multiplanar reconstructed image of the medullary tubule cavity at location A, i) a scanning electron microscopy (SEM) cross-sectional image from location C, and j) a magnified coronal multiplanar reconstructed image from location C.

The hoof wall is known to have a gradient in stiffness *in vivo* due to the inner material’s relatively higher exposure to hydration [44]. However, when tested in identical hydration conditions, samples from the outer hoof wall exhibited higher stiffness than those from the inner section [17]. Moreover, nanoindentation studies in the literature indicate that the hoof wall has a higher reduced modulus and hardness near the outer surface [45]. These results seemingly contradict that tubule density and area fraction porosity are higher near the stratum externum [15]. One would expect stiffness to decrease with increasing porosity. The tissue observed within the TMCs of tubules near the outer surface resolves this contradiction. By measuring the porosity with the area fraction method, the material appears to have higher porosity near the outer wall. Still, when the TMCs are observed with high-resolution μ-CT, more material within the tubules contributes to the overall stiffness.

Figure 6c shows that at location B, the tubules begin to have well-defined walls without any tissue within the TMC region. However, the coronal longitudinal multiplanar reconstructed view in Fig. 6f shows fine white lines within the TMC region, believed to be the beginnings of the bridges previously described by Huang *et al*. [9] and quantified by Lazarus *et al*. [24]. As we move toward the stratum internum, we observe an increasing number of bridges with thicker walls (Fig. 6d, g). The thin layer of a soft tissue-like structure within the TMC region in Fig. 6i appears to be a part of the bridge structure observed in Fig. 6j.

### 3.4 Helium pycnometer

The average porosity obtained from the helium pycnometer is approximately 8.07%, which is higher than the porosities reported by Huang *et al*. (3%) [10] and Lazarus *et al*. (0.77 ± 0.3%) [24]. This discrepancy is attributed to variations in sample location within the hoof and the analysis method used. The pycnometer employs helium gas, with helium molecules small enough to penetrate all types and sizes of pores, including the nanopores that cannot be detected by other imaging methods. Furthermore, the pycnometer measures the open porosity only, whereas other techniques measure the total porosity (open and closed porosities).

### 3.5 SBF-SEM

Figure 7 displays 2D images obtained using the SBF-SEM technique, showcasing the TMC region, tubule wall, and intertubular region. The white color in the images indicates the hollow area inside the tubules (TMC), which appears bright due to impregnation with epoxy resin. The darker spots represent the soft tissue enhanced by osmium and potassium ferrocyanide absorption. Based on Fig. 7, we conclude that the tubule wall (between the green lines) has nanoscale pores, penetrated by the resin. Leach [17] suggested that the horse hoof might have a similar permeability to the human nail since the horse hoof has a similar ultrastructure to the nail palate [46]. The porosity obtained from the helium pycnometer (Section 3.4) supports this phenomenon, as it is higher than the porosity measured using other techniques. The difference is due to nanopores within the tubule wall, as revealed by SBF-SEM.

**Fig. 7.**
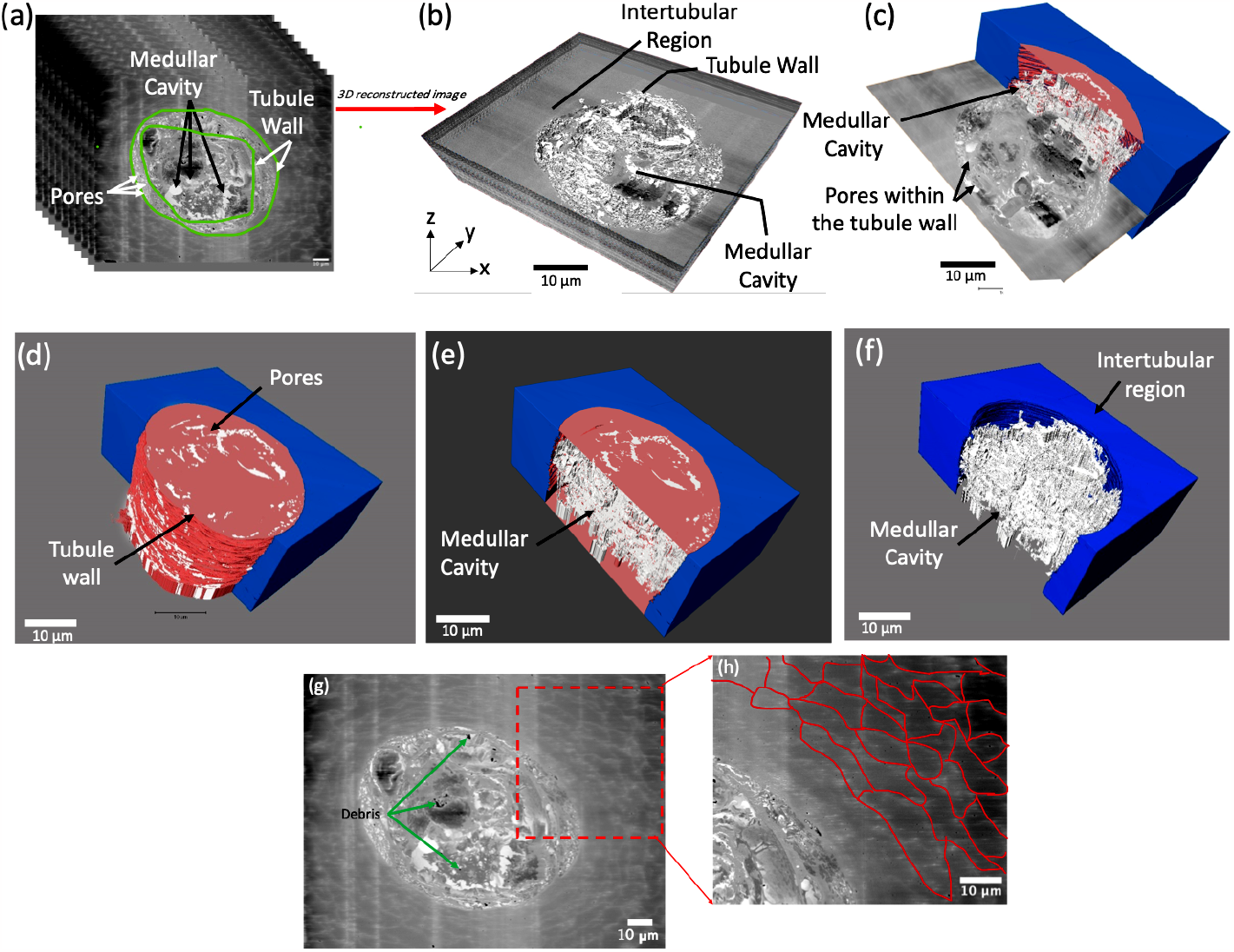
SBF-SEM images of a hoof wall specimen. (a) Stack of SBF-SEM scan images, (b) 3D volume rendering obtained by semi-automatic segmentation using Amira software, (c) 3D volume rendering displaying the inner and side walls of the tubule, (d) 3D volume rendering showing the TMC and intertubular area (shown in blue), (e) 3D volume rendering displaying a sagittal view of the TMC, (f) 3D extracted volume of the TMC with the tubule wall subtracted, (g) single slice from an SBF-SEM volume with green arrows indicating locations of debris on the sample surface during the slicing process, and (h) depiction of cell boundaries in the intertubular region highlighted in red.

Porous materials absorb energy in compression through the collapse of pores. During collapse, the force does not increase with increasing deformation [47, 48]. In the case of the hoof wall, collapsing pores could limit the maximum force transmitted to the internal structure of the hoof. Nanoscale pores observed from SBF-SEM would collapse and densify at very low strains, while the mesoscale tubules would collapse at higher strains [43]. Therefore, hierarchical porosity could be an essential structural feature for limiting the load transferred at different stages of impact.

Figures 7a, b, and c show that TMC is not entirely hollow, as previously reported [6, 17]. Figure 7d shows a 3D reconstructed image of a solitary tubule where the blue area represents the intertubular region and the white area depicts the hollow spaces within the TMC region, colored brown. Figure 7e presents a sagittal view of the tubule’s inner section, showing irregular tissue within the inner wall of the TMC instead of being entirely hollow. The hollow areas and the tubule’s inner wall were successfully extracted and represented in 3D in Fig. 7f using SBF-SEM and Amira software. The reconstructed image illustrates the irregular surface of the tubule’s inner wall. SBF-SEM imaging supported the results obtained earlier from the coronal multiplanar reconstructed image (Fig. 6), indicating that the TMC contains soft tissue and its inner wall is not smooth. A single 2D image from the stacked volume is illustrated in Fig. 7g, where the magnified area depicted by the overlaid irregular red line in Fig. 7h shows cell boundaries that appear as a wavy structure. The black particles shown on the sample’s surface in Fig. 7g resulted from falling debris generated during the cutting process inside, a common problem encountered in SBF-SEM [31].

## 4. Conclusions

We explored the hierarchical structure of the hoof wall. We evaluated various measurement methods for quantifying hoof wall porosity using state-of-the-art 3D multimodal imaging techniques, including μ-CT and SBF-SEM. Additionally, we applied the helium pycnometer technique, a novel approach within the field of biological materials characterization, to further enhance our understanding of the hoof wall porosity. We conducted μ-CT imaging at resolutions of 0.53 μm and 13.35 μm to obtain a comprehensive statistical analysis of the hoof wall microstructure and TMCs region. SBF-SEM was used for the first time to study the microstructure of the hoof wall at the nanoscale level. Our conclusions from this research are as follows:

1. 3D images acquired from μ-CT indicated that the tubules have thin wall bridges segmenting them into air pockets, which start in the middle section of the hoof wall and increase in density towards the inside of the hoof wall.
2. The tubules exhibit some degree of periodic waviness, which may be an intrinsic feature or a result of dryness and moisture loss caused by X-ray exposure during μ-CT testing.
3. High-resolution μ-CT revealed tubules are filled with tissue near the outer surface and empty in the middle to inside sections of the hoof wall.
4. Statistical analysis of the TMCs indicated that the average values of Eqdiameter, surface area, volume, and sphericity are 69.43 μm, 3.55 ×10^4^ μm^2^, 4.53 ×10^5^ μm^3^, and 0.86, respectively.
5. Dividing the hoof wall into different zones showed that the porosity increases from the inner to the outer direction of the hoof wall thickness.
6. Different techniques yielded different porosity measurements. While μ-CT 2D reported similar values of 2.98% to previous studies, the 3D calculation showed an increase in porosity to 3.47%, which is considered more accurate because the estimates are based on the voxel size of the pores, not the cavity’s radius.
7. Helium pycnometer measurements detected nanoscale pores that were not visible using standard imaging techniques like μ-CT or SEM, which resulted in a higher average porosity of 8.07%. Thus, it is an accurate technique that can detect pore sizes ranging from microscale to nanoscale.
8. SBF-SEM successfully provided a nanoscale image of the hoof wall, which is challenging to obtain using transmission electron microscopy or any 3D microscopy technique. SBF-SEM imaging showed nano-sized pores within the tubule wall, which have not been previously detected, and explains the higher porosity observed using the helium pycnometer.

Mechanics studies of the hoof wall and bioinspired materials under impact will benefit from the morphometric data presented in this work. Drop tower tests indicated tubule density strongly affects crack propagation in bioinspired samples [24], and the elliptical shape and bridges of tubules are hypothesized to have essential mechanical functions [17, 24]. Therefore, it is crucial to identify methods to accurately quantify the structure of the hoof wall with metrics such as sphericity, porosity, and tubule density. Moreover, the hierarchical porosity observed in this work should be further explored as a potential design motif for bioinspired impact-resistant materials.

The present study has some limitations. The number of samples is limited to specific locations and cannot be generalized to other areas within the hoof wall. Additionally, the sample was extracted from one horse, representing only one species and gender.

## Declaration of Generative AI and AI-assisted technologies in the writing process

During the preparation of this work, the authors used GPT-3.5 to improve readability and clarity. After using this tool, the authors reviewed and edited the content as needed and take full responsibility for the content of the publication.

## Acknowledgments

The authors acknowledge the researchers at the Core facilities at the Carl R. Woese Institute for Genomic Biology (IGB) for their assistance with micro-computed tomography (μ-CT) and serial block-face scanning electron microscopy (SBF-SEM). The staff scientists of the Microscopy Suite at the Beckman Institute for Advanced Science and Technology at the University of Illinois at Urbana-Champaign are also acknowledged for their help in performing μ-CT imaging. The authors would also thank Ahmed Adel Hassan, a Ph.D. candidate at the Department of Civil Engineering at the University of Illinois at Urbana-Champaign, for developing the 3D visualization of the hierarchical structure of the hoof wall shown in Fig. 1. Lastly, the authors gratefully acknowledge the National Science Foundation (NSF) for the support received under Grants No. MOMS-1926353 and 1926361.

## A.1 Image processing for μ-CT

Micro-CT or μ-CT is a non-destructive 3D imaging technique to obtain the internal structure of an object. In this technique, x-rays generated using an X-ray source are transmitted through the samples. An X-ray detector then records the X-rays as a 2D projection image. The sample is then rotated, and the same procedure is followed. The projection (imaging) is continuously collected while the sample is rotating continuously at a fixed angular rotational angle while repeatedly irradiated with an X-ray beam. Upon the interaction between the X-ray beam and the sample, the beam gets attenuated (loses its initial energy) as it passes through the sample by the sample matrix [48]. X-ray beam attenuation is a function of beam energy (characterized by intensity and flux density), which is controlled by the atomic number and density of the object being scanned [48]. Lambert Beer’s law describes monochromatic X-ray beam attenuation by demonstrating the transmitted intensity I of a beam as it propagates through an object in a straight pass:

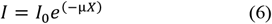

where ***I***_**0**_ is the integral current of the X-ray photon (initial photon intensity), ***I*** is the integral current transmitted by the sample (final photon intensity), μ is the linear attenuation coefficient of the specimen being scanned [L-1], and X is the specimen thickness. The linear attenuation coefficient (μ) value depends on the bulk density of the sample, photon energy, and the electron density of the scanned materials [49].

The 2D images produced from the scan are combined and reconstructed to develop 3D images using the 3D volume rendering procedure, which provides an excellent representation of the internal structure of the investigated materials. Therefore, μ-CT is considered a powerful non-destructive technique for analyzing the variation in the densities and visualizing the internal structure of most materials. A more detailed discussion about X-ray computer tomography principles is discussed intensively by [42, 50-52].

Quantitative analyses of μ-CT data are considered the primary purpose of image processing which is done by reconstructing and visualizing sample volume via 3D rendering procedures. Non-destructive 3D reconstruction techniques are used herein to obtain a comprehensive statistical analysis of the hoof wall. Various software such as (Avizo, pore 3D, VSG, etc.) is now commercially available for textural and morphological quantification of the internal constituents of different materials. The quantitative analysis consists of multiple steps, which start with a volume of interest (VOI) selection. In our case, the tubule area is considered VOI. The image segmentation process is done by dividing the image volume into different regions represented by a group of voxels belonging to the same phase.

The segmentation algorithms are divided into two categories determined by the discontinuity and similarity of the beam intensity values [53]. The intensity discontinuity category is by partitioning the image based on the sudden change in the intensity. In contrast, the similarity intensity category is partitioning the image based on areas with similar intensities using the thresholding method [53]. There are many thresholding methods, including global threshold, basic adaptive threshold, clustering, and region-growing methods. The choice of the efficient thresholding technique is based on the degree of the morphological complexity of the required study area in the sample. For example, geometry in the pore spaces is more complex since they have irregular shapes and are at times filled with mixed phases (air, liquid, solid), making it difficult to separate them efficiently using the automatic thresholding method (global thresholding). Therefore, more sophisticated methods such as refined thresholding techniques, basic adaptive thresholding, and clustering by iterative methods are more efficient in such cases. However, one threshold value used in the global threshold method is enough in portioning the image when the gray-level histogram is bimodal or multimodal but cannot be applied in unimodal histogram images [53, 54]. The last step is refining the image from unwanted pixels containing more than one phase caused by artifacts such as the partial volume effect [42].

The goal of segmentation in our case was to isolate the TMC region from the hoof wall intratubular region to visualize and perform statistical quantification for the tubule’s volume, area, sphericity, and equivalent diameter (Eqdiameter) in 3D. For better accuracy for the segmentation and to ensure that no incomplete tubule was included in the analysis, the segmentation process starts with selecting a cubical volume of interest in the middle of each specimen which is away from the concave corners of the hoof sample. Selecting a threshold value is crucial for a successful segmentation procedure. A Series of known processing steps for tubule segmentation following

Avizo software [55] was performed. Initially, interactive threshold modules were used for tubule detection to produce a binary image where the intensity level is 1 for the cavity, and the 0-intensity level is for the hoof wall intratubular region. A morphological opening operator was applied to remove all the small objects and smoothen the boundaries of the interesting feature (tubules medullary cavities), producing filtered images free of artifacts and noise. Finally, a labeling analysis module was used to perform a set of computational measurements for each particle in a 3D image.

To determine the area fraction porosity through the hoof wall thickness, images were analyzed using Fiji [56] software. Two images are selected for analysis: one on the top and one on the bottom of each hoof wall sample. The images are cropped such that the sections to analyze are tangential to the Stratum Internum, and further cropped into smaller sections. The threshold, that was manually applied, ranged between 2-7% between the four hoof wall samples to include all microtubules while eliminating the background noise. A median filter, DE speckle, was applied to further eliminate the background noise. The threshold and median filter parameters were applied consistently for each hoof wall sample. The command, Particle Analyzer, measured the tubule areas and total area of the crop section.

## A.2 Image processing for SBF-SEM

The data collected using SBF-SEM does not require further alignment because the images are acquired continuously during the scan before sectioning. Therefore, the collected images can be staked in volume files using software such as AmiraTM [32]. 3D Serial Block Face SEM datasets were batch-converted into a total of 1426 tiff files in preparation for modeling in Amira™ (FEI), following the method described by [57] with a few modifications. About 1426 data stacks were imported into Amira™, and the appropriate voxel dimensions were inputted when prompted. Semi-automated segmentation was performed using a combination of the threshold, the magic wand, the lasso, and interpolation tools, allowing parameters such as volume and surface area to be measured. The magic wand, an automatic tool that uses a polygon expansion based on a voxel contrast gradient to select a connected group of triangles, was used to segment the pores. The lasso and interpolation tools were used to segment the hoof walls. The rest of the block was segmented using the threshold tool. Each segmented material was statistically analyzed using the label analysis tools and filtered out using the threshold by criterion and filter by measure tools. The volume rendering and generated surface tools were then used to render the segmented data in 3D.

